# pTDP-43 aggregates accumulate in the gut and other non-central nervous system tissues prior to symptom onset in amyotrophic lateral sclerosis

**DOI:** 10.1101/2022.03.17.484805

**Authors:** Samuel B. Pattle, Judi O’Shaughnessy, Olivia M. Rifai, Judith Pate, Mark J. Arends, Fergal M. Waldron, Jenna M. Gregory

## Abstract

**Objective:** Neurodegenerative diseases such as Parkinson’s disease (PD), Alzheimer’s disease (AD) and amyotrophic lateral sclerosis (ALS) are traditionally considered strictly neurological disorders. However, clinical presentation is not restricted to neurological systems, and non-central nervous system (CNS) manifestations, particularly gastrointestinal (GI) symptoms, are common. Our objective was to understand the systemic distribution of pathology in archived non-CNS tissues, taken as part of routine clinical practice during life from people with ALS.

**Design:** We requested all surgical specimens of non-CNS tissue taken during life from 48 people with ALS, for whom evidence of the characteristic proteinopathy associated with ALS had been identified in the CNS after death (i.e., the pathological cytoplasmic accumulation of phosphorylated TDP-43 (pTDP-43) aggregates). Of the 48 patients, 13 had sufficient tissue for evaluation: 12 patients with sporadic ALS and 1 patient with a *C9orf72* hexanucleotide repeat expansion. The final cohort consisted of 68 formalin-fixed paraffin embedded tissue samples from 22 surgical cases (some patients having more than one case over their lifetimes), representing 8 organ systems, which we examined for evidence of pTDP-43 pathology. The median age of tissue removal was 62.4 years old and median tissue removal to death was 6.3 years.

**Results:** We identified pTDP-43 aggregates in multiple cell types of the GI tract (i.e., colon and gallbladder), including macrophages and dendritic cells within the lamina propria, as well as neuronal and glial cells of the myenteric plexus. Aggregates were also noted within lymph node parenchyma, blood vessel endothelial cells, and chondrocytes. We note that in all cases with non-CNS pTDP-43 pathology, aggregates were present prior to ALS diagnosis (median=3years) and, in some instances, preceded neurological symptom onset by more than 10 years.

**Conclusion:** These data imply that patients with non-CNS symptoms may have occult protein aggregation that could be detected many years prior to neurological involvement.

**Summary:** Neurodegenerative diseases such as Parkinson’s disease (PD), Alzheimer’s disease (AD) and amyotrophic lateral sclerosis (ALS) are traditionally considered strictly neurological disorders. However, clinical presentation is not restricted to neurological systems, and non-central nervous system (CNS) manifestations, particularly gastrointestinal (GI) symptoms, are common. Our objective was to understand the systemic distribution of TDP-43 pathology in archived non-CNS tissues, taken as part of routine clinical practice during life from people with ALS. We identified pTDP-43 aggregates in multiple cell types of the GI tract (i.e., colon and gallbladder), and within lymph node parenchyma, blood vessel endothelial cells, and chondrocytes. We note that in all cases with non-CNS pTDP-43 pathology, aggregates were present prior to ALS diagnosis (median=24months) and, in some instances, preceded neurological symptom onset by more than 10years. These data imply that patients with non-CNS symptoms may have occult protein aggregation tha could be detected many years prior to neurological involvement.

**Graphical Abstract:** Graphical Abstract.
Ante-mortem tissue cohort comprised of tissue taken from people with ALS demonstrates non-CNS accumulation of pTDP-43 aggregates prior to symptom onset.
Schematic of workflow to identify pTDP-43 aggregates indicative of non-central nervous system (CNS) manifestations of ALS. Lower panel left: cartoon depicting organs and cell types that had evidence of pTDP-43 aggregation in ALS patient non-CNS ante-mortem tissue. Lower panel right: cartoon depicting organs with no evidence of pTDP-43 aggregation in ALS patient non-CNS ante-mortem tissue.

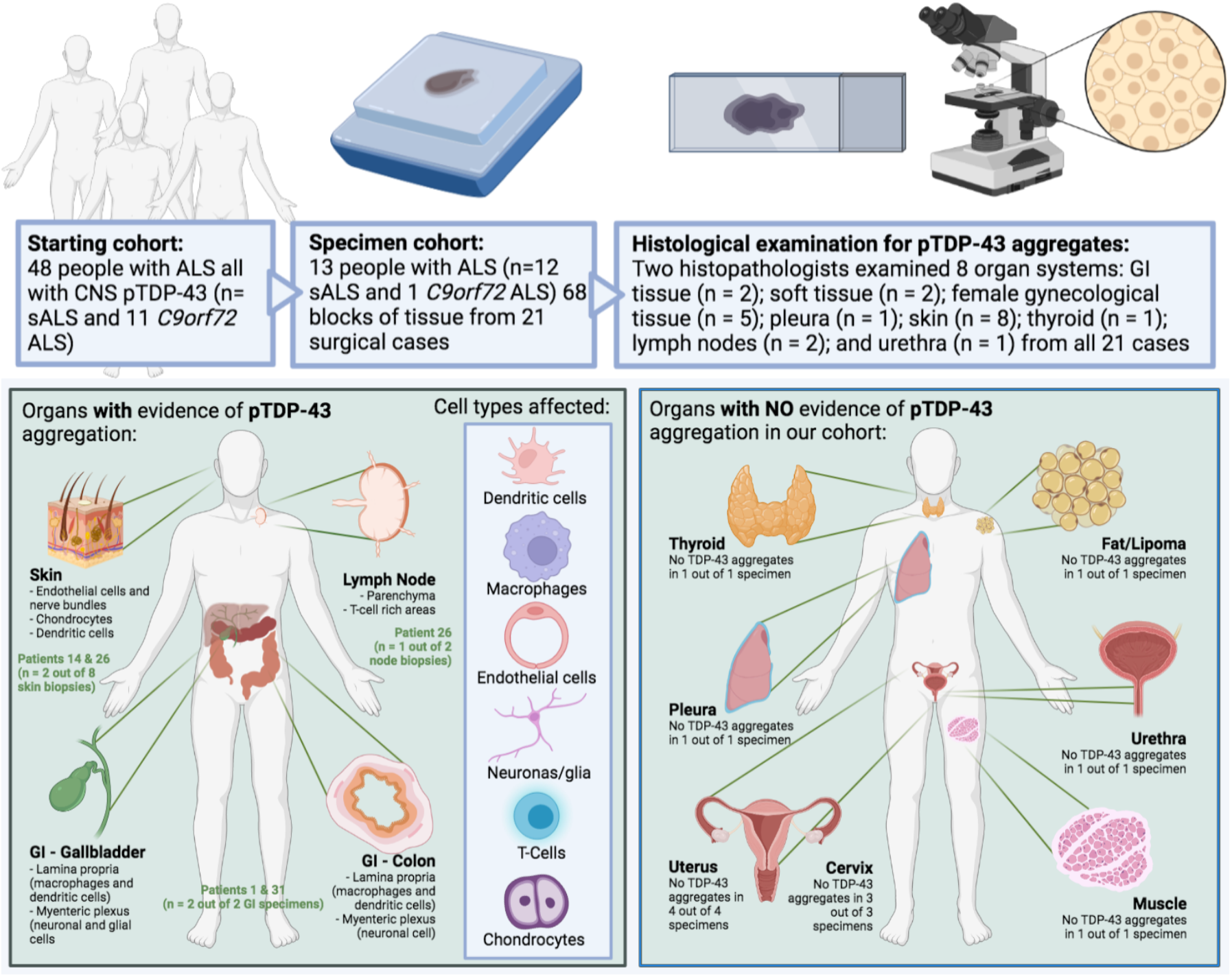

## Introduction

Neurodegenerative diseases such as Parkinson’s disease (PD), Alzheimer’s disease (AD), and amyotrophic lateral sclerosis (ALS) are traditionally classified and managed as neurological disorders. There is, however, marked clinical heterogeneity within these disorders, which may in part contribute to the large number of failed clinical trials, and clinical presentation is not restricted to neurological dysfunction [1]. Indeed, non-central nervous system (CNS) manifestations, including gastro-intestinal (GI) symptoms (e.g. weight loss and altered bowel habit), are common in these patients [1]. Importantly, to date, biomarker and therapeutic studies in the neurodegeneration field have been primarily aimed at assessing pathways known to be perturbed within the CNS, and in a clinically diverse patient population. It should be possible to broaden the therapeutic potential for people with neurodegenerative diseases by expanding our approach to include clinical trials targeted at non-motor symptoms, with the aim of improving quality of life and even reducing the non-CNS symptom burden for people with ALS. This approach demands a deeper understanding of the systemic distribution and molecular drivers of these non-CNS symptoms.

Systemic non-CNS manifestations of PD and other neurodegenerative diseases are now well-established [2,3]. Furthermore, a population-wide study of almost 1 million people in a Swedish cohort between 1965 and 2016 demonstrated that individuals with GI symptoms who underwent a colonoscopy with biopsies that were reported as histologically normal were statistically at a higher risk of developing ALS (Hazard ratio = 1.22; 95% confidence interval = 0.94-1.51) [4]. These data imply that patients with GI symptoms could have occult protein aggregation that could be detected diagnostically many years prior to neurological involvement. Indeed, alpha-synuclein has been shown to accumulate in pathological aggregates in the GI tract of PD patients up to 20 years before CNS manifestations begin [5], raising the possibility that such biomarkers may also hold promise as early indicators of disease.

The unifying proteinopathy seen in all cases of sporadic ALS and most genetic ALS cases is the cytoplasmic accumulation of phosphorylated TDP-43 (pTDP-43) aggregates in the central nervous system [6]. Several studies have demonstrated pTDP-43 aggregates within muscle and peripheral nerve biopsies from patients with ALS, using western blotting, immunohistochemistry, and electron microscopy [7-9]. Furthermore, several TDP-43 animal models published to date have demonstrated GI pathology that occurs prior to neurological symptom onset [10-12]. However, no study to date has evaluated human GI tissue or other non-CNS tissues in a systematic way in this patient group. Here we examine non-CNS tissues, including GI specimens, from a cohort of people with ALS to assess the systemic distribution and burden of TDP-43 pathology in archived tissues, taken as part of routine clinical practice during life from people with ALS.

## Methods

### Human tissue acquisition and ethics

All tissue was requested from the Lothian NRS BioResource RTB ethical approval 15/ES/0094, covering use of residual tissue surplus to diagnostic requirements taken as standard of care with approval number SR1684. All archived formalin-fixed, paraffin-embedded (FFPE) tissue material with an NHS Lothian diagnostic code (e.g. any tissue samples taken for diagnostic or surgical purposes at any time during the patient’s life, under the care of NHS Lothian and NHS Fife) was requested for 48 patients with known diagnosis of sporadic ALS or ALS with a *C9orf72* hexanucleotide repeat expansion. The tissue was then assessed for sufficiency by two independent histopathologists (SBP & JMG) using a single H&E. All clinical data, in addition to data provided by the Lothian NRS BioResource were non-identifiable and collected as part of Scottish Motor Neurone Disease Register (SMNDR) and Care Audit Research and Evaluation for Motor Neurone Disease (CARE-MND) platform (ethics approval from Scotland A Research Ethics Committee 10/MRE00/78 and 15/SS/0216) and all patients consented to the use of their data during life. All demographic and clinical data are collated in Table 1. Tissue provided by the BioResource had been fixed in 10% formalin for a minimum of 72 h, then dehydrated in an ascending alcohol series (70–100%) followed by three 4 hour washes in xylene. Three successive 5-hour paraffin wax embedding stages were performed followed by cooling and sectioning of tissue on a Leica microtome into 4 × 4 μm thick serial sections that were collected on Superfrost (ThermoFisher Scientific) microscope slides. Sections were dried overnight at 40°C before immunostaining.

**Table 1.** Demographic and pathological data for the 13 ALS patients comprising the ante-mortem tissue cohort examined. Clinical demographics and summary of tissue type, diagnosis and findings from pTDP-43 staining for the tissue cohort comprising the 13 ALS patients with examinable material.

| Patient ID | Cohort | N° tissue blocks | Age at sampling | Age at death | Specimen type | Diagnosis | pTDP-43 pathology | Cell types / regions affected | If TDP-43 present - time to: Symptom onset | ALS diagnosis |
| --- | --- | --- | --- | --- | --- | --- | --- | --- | --- | --- |
| 1 | sALS | 2 | 62 | 65 | Colon | Normal | Present | Myenteric plexus and lamina propria | Coincident | 2 months |
|  |  | 6 | 62 |  | Muscle | Denervation | Present |  |  |  |
| 3 | sALS | 6 | 53 | 72 | Thyroid | Goiter (adenomatous) | Absent | Endothelial cells |  |  |
| 11 | sALS | 1 | 45 | 59 | Endometrium | Endometrium (secretory) | Absent |  |  |  |
|  |  | 1 | 57 |  | Endometrium | Endometrium (inactive) | Absent |  |  |  |
| 14 | sALS | 1 | 54 | 61 | Skin | Seborrheic keratosis | Present | Dendritic cells; endothelial cells; nerve bundles | 48 months | 60 months |
| 17 | sALS | 1 | 44 | 56 | Cervix | Polyp | Absent |  |  |  |
|  |  | 1 | 44 |  | Skin | Seborrheic keratosis | Absent |  |  |  |
|  |  | 6 | 46 |  | Uterus | Leiomyoma | Absent |  |  |  |
| 18 | sALS | 2 | 73 | 74 | Urethra | Chronic inflammation | Absent |  |  |  |
| 26 | sALS | 3 | 68 | 70 | Skin | Basal cell carcinoma | Present | Chondrocytes; endothelial cells; dendritic cells; nerve bundles in skin. Interfollicular/paracortical lymph node parenchyma (T-cell rich areas of node) | Coincident | 12 months |
|  |  | 5 | 55 |  | Lipoma and lymph node | No malignancy | Present |  | 144 months | 168 months |
| 28 | sALS | 2 | 74 | 76 | Lymph node | Unsatisfactory | Absent |  |  |  |
|  |  | 3 | 74 |  | Lymph node | No malignancy | Absent |  |  |  |
|  |  | 3 | 70 |  | Pleura | No malignancy | Absent |  |  |  |
| 29 | C9orf72 | 1 | 55 | 66 | Skin | Squamous cell ca in situ | Absent |  |  |  |
|  |  | 1 | 57 |  | Skin | Hyperkeratosis | Absent |  |  |  |
| 30 | sALS | 2 | 67 | 73 | Skin | Pigmented nevus | Absent |  |  |  |
|  |  | 12 | 61 |  | Uterus | Adenocarcinoma | Absent |  |  |  |
| 31 | sALS | 2 | 75 | 78 | Gall Bladder | Gallstone disease | Present | Myenteric plexus and lamina propria | 12 months | 24 months |
| 33 | sALS | 1 | 83 | 84 | Skin | Squamous cell carcinoma | Absent |  |  |  |
| 40 | sALS | 2 | 71 | 74 | Skin | Actinic keratosis | Absent |  |  |  |

### H&E staining

Sections were dewaxed using successive xylene washes followed by alcohol hydration. H&E staining was performed by first immersing slides in Harris Haematoxylin for 5 minutes. Following this, slides were dipped twice into 1% acid alcohol and immediately washed in running tap water. Slides were then immersed in lithium carbonate until they turned blue and then washed again in running tap water. Slides were then dehydrated by agitating the rack and slides in alcohol and then immersed in eosin Y for 2 minutes followed by 3 successive washes in 100% alcohol for 30 seconds. The slides were then cleared in two changes of xylene then mounted using DPX mountant. Sections were imaged using an Olympus BX51 with a Nikon Digital-Sight camera.

### Immunohistochemistry

Sections were dewaxed using successive xylene washes followed by alcohol hydration and treatment with picric acid for removal of formalin pigment. For pTDP43 and CD68 staining, antigen retrieval was carried out in citric acid buffer (pH 6) in a pressure cooker for 30 min, after which immunostaining was performed using the Novolink Polymer detection system with a 2B Scientific (Oxfordshire, UK) anti-phospho(409–410)-TDP43 antibody at a 1 in 4000 dilution, Agilent (Cheadle, UK) anti-CD68 antibody at a 1 in 100 dilution. Tau immunostaining required no antigen retrieval and the Phospho-Tau (Ser202, Thr205) Monoclonal Antibody (AT8) antibody was used at a 1 in 2000 dilution (ThermoFischer, UK). Immunodetection was performed using 3,3’-diaminobenzidine (DAB) chromogen and slides were counterstained with haematoxylin and lithium carbonate and mounted using DPX mountant. Sections were imaged using an Olympus BX51 with a Nikon Digital-Sight camera. Bleach treatment was performed (on selected cases where warranted) by incubating slides immediately post antigen retrieval in 10% hydrogen peroxide for 12 hours overnight. This step was followed by the normal protocol in full, including the additional 3% hydrogen peroxide step used to block endogenous peroxidase activity, following a previously published and optimised protocol [13].

## Results

### Establishment of a cohort of ALS non-CNS tissue

We have previously evaluated pTDP-43 staining within the CNS of 48 patients (37 sporadic ALS patients and 11 patients with a *C9orf72* hexanucleotide repeat expansion). All 48 patients had pathological pTDP-43 inclusions in their CNS tissue and a diagnosis of ALS, and their CNS *post-mortem* tissue has been evaluated previously in our prior studies [14,15]. Within this group of 48 individuals with a diagnosis of ALS, 9 had mild cognitive involvement (ALDci; measured by the Edinburgh Cognitive ALS Screening (ECAS) tool) and one had a behavioural deficit (ALSbi), but none met criteria for a diagnosis of frontotemporal dementia (FTD) and 32 of the cases had not been evaluated using the ECAS during life. All cases included in the cohort had CNS pTDP-43 pathology in motor brain regions and in some cases, also in non-motor brain regions [14], and we wanted to establish whether they also had pTDP-43 pathology within non-CNS tissue. As such, we requested all non-CNS tissue from the NHS Lothian BioResource, a local tissue bank that provides controlled access to all tissue taken during life (i.e., diagnostic and surgical) from all patients under the care of NHS Lothian and Fife, for approved research use. Of the 48 patients that we requested tissue for, 13 had surgical specimens from non-CNS sites and sufficient tissue for evaluation: 12 patients with sporadic ALS and 1 patient with a *C9orf72* hexanucleotide repeat expansion. The final cohort consisted of tissue from 22 surgical cases (some patients having more than one case over their lifetimes), representing 8 organ systems: GI tissue (n = 2), soft tissue (e.g. lipoma/muscle; n = 2), female gynecological tissue (n = 5), pleura (n = 1), skin (n = 8), thyroid (n = 1), lymph nodes (n = 2), and urethra (n = 1) (Table 1). Interestingly, pTDP-43 aggregates were only identified in GI tissue (2 out of 2 cases), lymph nodes (1 out of 3 cases) and skin (2 out of 8 cases). For those with aggregates, the median time to symptom onset and diagnosis from tissue sampling was 12 months (range = 0-144 months) and 24 months (range = 2-168 months), respectively. Both GI specimens evaluated were taken 2 months and 2 years prior to diagnosis, with the gallbladder specimen showing pTDP-43 aggregates 1 year prior to the onset of their motor symptoms. Notably, one sample from patient 26 had pTDP-43 aggregates present, in a lymph node, 14 years prior to diagnosis with ALS. The remaining tissues showed no evidence of pTDP-43 pathology other than within blood vessel endothelial cells (in 6 out of 22 specimens; Table 1).

### Presence of pathological pTDP-43 aggregates in gastrointestinal tract of ALS patients prior to motor symptom onset highlights potential for use as early biomarker

Patient 1 (Table 1) had an investigative colonoscopy around the time of ALS symptom onset, 2 months prior to their ALS diagnosis, and the examination revealed a small sessile polyp; three random colonic biopsies were also taken at the time of biopsy. The sessile polyp was a benign, non-dysplastic hyperplastic polyp and the three random colonic biopsies were reported as normal with no evidence of infection or microscopic colitis and no evidence of dysplasia or malignancy. Both the polyp and the colonic biopsies showed evidence of pTDP-43 aggregation within the lamina propria (the mucosal connective tissue deep to the surface enterocytes; Figure 1A) with examples of pTDP-43 aggregates in CD68-positive (i.e., activated) macrophages (Figure 1B) and in non-CD68-positive dendritic cells (Figure 1C). pTDP-43 aggregates were also visualized within neuronal cells in the nerve bundle sampled in one of the colonic biopsies (Figure 1D). The pTDP-43 aggregates were seen to have multiple morphologies, including single dense perinuclear amorphous cytoplasmic aggregates (Figure 1E; top panel) and scattered dispersed small dot-like cytoplasmic staining (Figure 1E; bottom panel), similar to that seen in the CNS. The aggregation in these specimens was specific for TDP-43 proteinopathy; no pTau aggregates were observed (Figure 1A).

**Figure 1.**
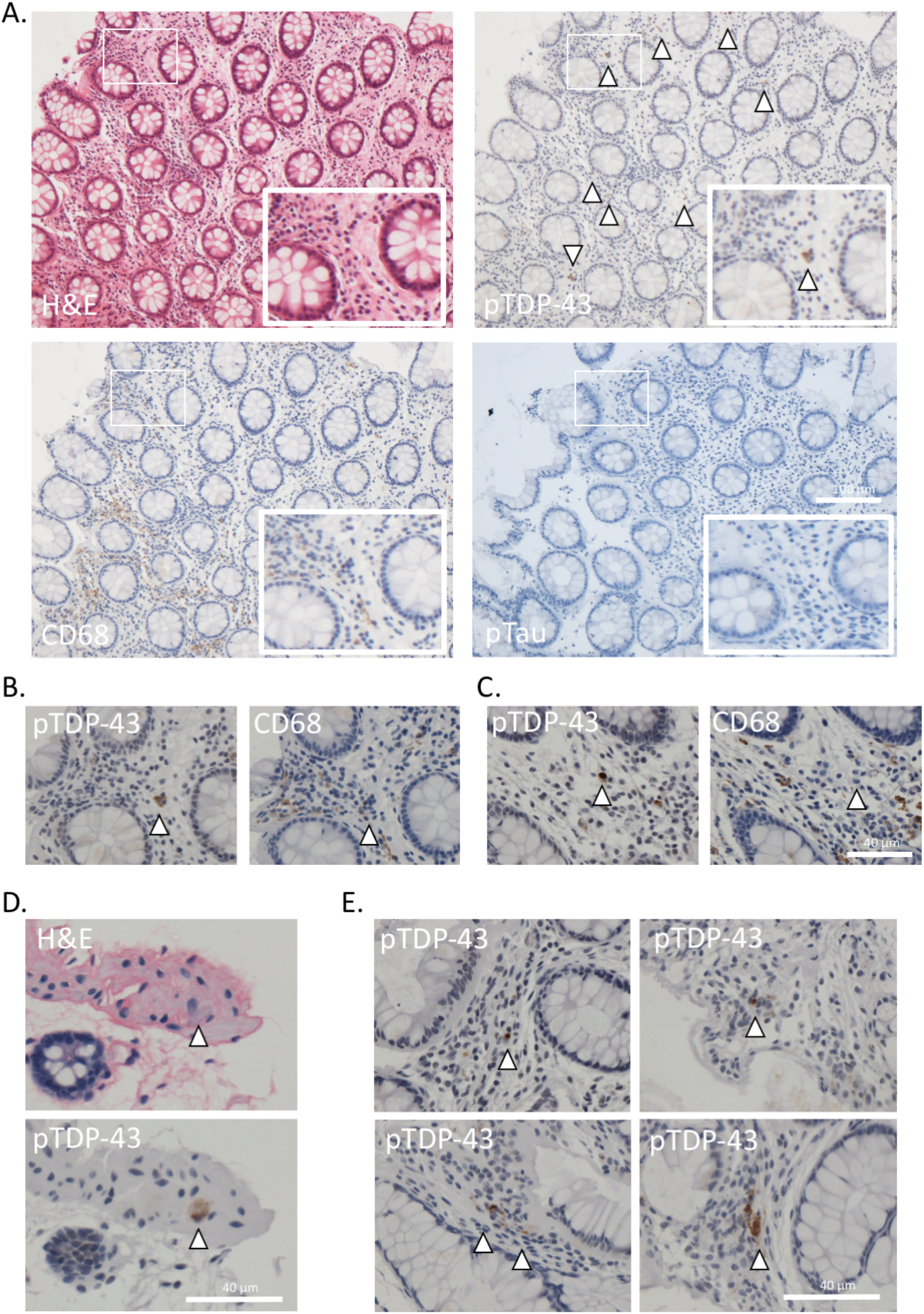
pTDP-43 aggregates are present in lamina propria and nerve bundles of colonic tissue and demonstrate diverse aggregate morphologies. **A**. Photomicrographs of colonic biopsy (patient 1). Top left H&E top right pTDP-43 (multiple lamina propria aggregates highlighted by white arrow heads); bottom left CD68 (highlighting activated macrophages); bottom right pTau (negative). Scale bar = 50 µm, images taken at 20x magnification. **B**. 40x magnification photomicrograph highlighting, in serial sections, macrophage staining in the same area as the TDP-43 aggregate staining (i.e. pTDP-43 present within macrophages, implying active aggregate clearance). **C**. 40x magnification photomicrograph highlighting, in serial sections, no evidence of macrophage staining in the same area as the cell, highlighted with a white arrowhead, containing a dense perinuclear single pTDP-43 aggregate within a dendritic cell (implying that pTDP-43 aggregates are also in cell types that are not clearing aggregates, i.e. non-macrophage cells). **D**. H&E (top) and pTDP-43 immunohistochemistry (bottom) images demonstrating pTDP-43 aggregation in a large neuronal cell within a nerve bundle. **E**. Examples of pTDP-43 aggregate morphology ranging from single, dense perinuclear aggregates (top images) and dispersed cytoplasmic dot-like aggregates throughout the cytoplasm (bottom images).

Patient 31 (Table 1) had a cholecystectomy 12 months prior to ALS symptom onset and 24 months prior to their ALS diagnosis, and the gallbladder was reported as showing chronic cholecystitis secondary to gallstones. pTDP-43 staining revealed extensive aggregate formation in neuronal and glial cells within the myenteric plexus (the primary neurovascular bundle of the submucosal tissue) of the gallbladder (Figure 2A-B). Aggregates were also noted within the endothelial cells of small-to medium-sized blood vessels within the myenteric plexus (Figure 2A). There was also evidence of extensive pTDP-43 aggregate formation within dendritic cells and macrophages in the lamina propria, as seen in the colonic mucosa (Figure 2C).

**Figure 2.**
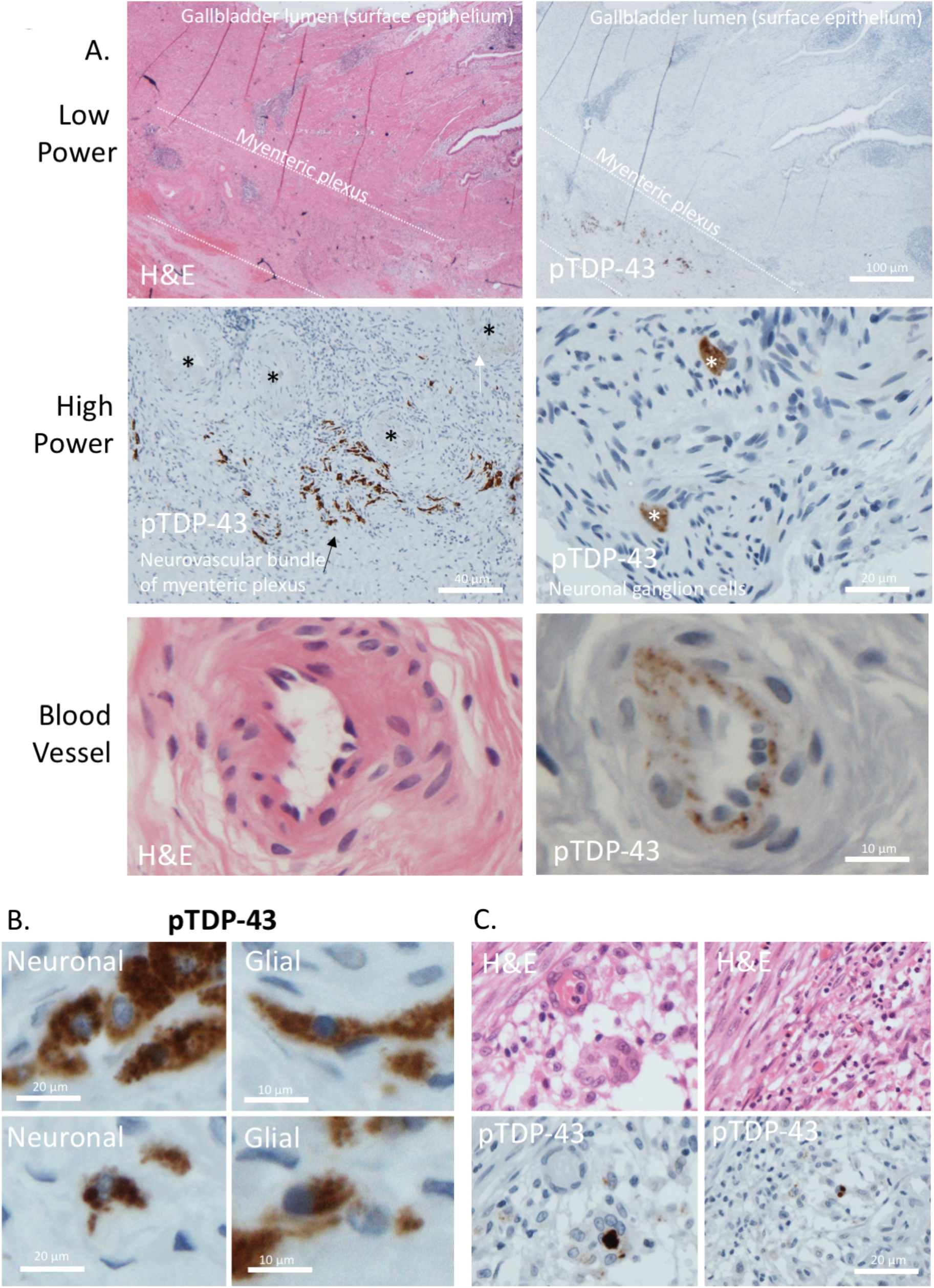
pTDP-43 aggregates identified in lamina propria and myenteric plexus of gallbladder mucosa. **A**. Low power photomicrographs of gallbladder wall (patient 31), top left -H&E illustrating location of epithelial-lined mucosal surface and myenteric plexus. Top right – pTDP-43 staining performed on the serial section showing extensive pTDP-43 aggregation within the myenteric plexus. Middle panel (left) high power view of myenteric plexus, with aggregates in the neuronal cells (black arrow) and endothelial cells (white arrow) of the plexus. Black asterisks show blood vessels cut in transverse section. Middle panel (right) two neuronal cells demonstrating cytoplasmic pTDP-43 aggregation within a nerve bundle in the wall of the gallbladder. Lower panel (left) H&E and (right) pTDP-43 staining demonstrating blood vessel endothelial cells cytoplasmic aggregates. **B**. Example photomicrographs demonstrating aggregates in cells with neuronal morphology (left) and glial morphology (right) **C**. Left, and right demonstrate H&E (top) and pTDP-43 performed on the serial section (bottom), demonstrating extensive lamina propria and mucosal aggregates within dendritic cells and macrophages.

### Pathological pTDP-43 aggregates are identified in lymph node parenchyma and specialized cells of the skin

Patient 26 had several specimens taken at different times, several years apart, all prior to their diagnosis of ALS. The first specimen of note was a probable lipoma taken along with a supraclavicular lymph node 14 years prior to their ALS diagnosis and 12 years before they had ALS-associated motor symptoms. The specimen, at the time of sampling, was reported as a lipoma and normal lymph node with no evidence of malignancy. Our examination revealed the lymph node to contain cells that stained positively for pTDP-43 (Figure 3). These cells appeared within the interfollicular/paracortical lymph node parenchyma, predominantly in T-cell-rich and antigen-presenting regions (Figure 3A). pTDP-43 aggregates were also found within blood vessels in the paranodal tissue (Figure 3B). Of note, no pTDP-43 aggregates were found in the muscle biopsy sampled at time of symptom onset, 2 months prior to their ALS diagnosis (Figure 3C). The next specimen of note was a skin sample taken from the ear around the time of their symptom onset, 12 months prior to their ALS diagnosis, with a clinical suspicion of basal cell carcinoma, and was reported as a completely excised basal cell carcinoma. The specimen showed evidence of pTDP-43 aggregates within several cell types and regions: (i) the superficial dermis of the skin (Figure 4A); (ii) within endothelial cells of small to medium sized blood vessels; (iii) within nerve bundles; and (iv) within chondrocytes (cartilage-producing mesenchymal cells which may have a shared lineage with neurons in the neural crest) (Figure 4B). The superficial dermis, neurons and endothelial cells were also seen to be involved by pathological pTDP-43 aggregates in tissues sampled from other cases (see Table 1 and graphical abstract for a summary of affected tissues).

**Figure 3.**
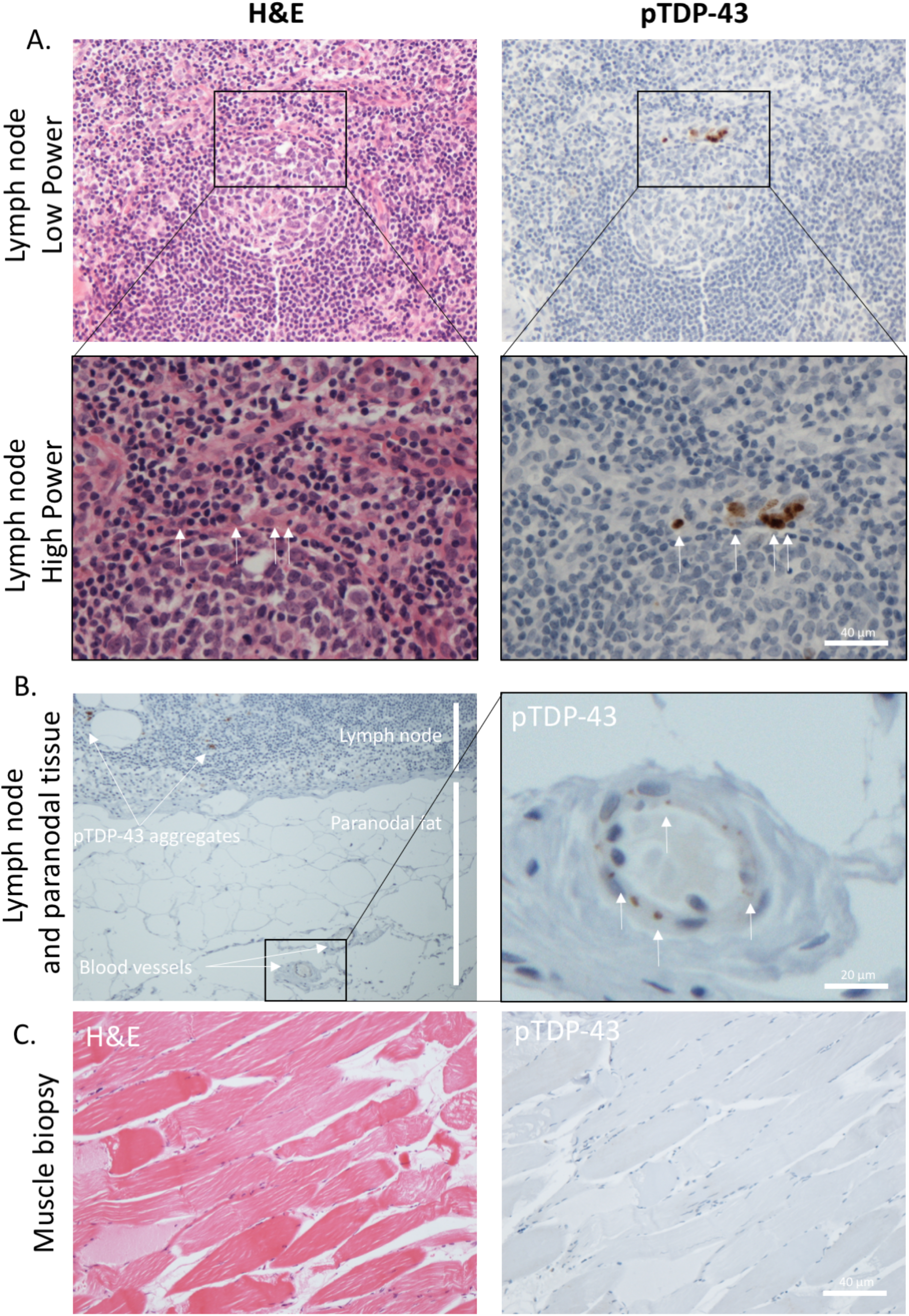
pTDP-43 aggregates identified in lymph node parenchyma, endothelial cells and chondrocytes long before symptom onset of ALS. **A**. H&E (left) and pTDP-43 (right), low power (top) and high power (bottom) images of an active lymph node germinal centre with evidence of cells containing pTDP-43 aggregates (white arrows) within the mantle zone rim of the lymphoid follicle. **B**. Low power (left) and high power (right) images of the paranodal tissue demonstrating blood vessel pTDP-43 aggregation (white arrows) in adjacent feeder vessels. **C**. H&E (left) and pTDP-43 (right) images showing no evidence of pTDP-43 aggregates in a muscle biopsy taken at point of diagnosis from patient 1.

**Figure 4.**
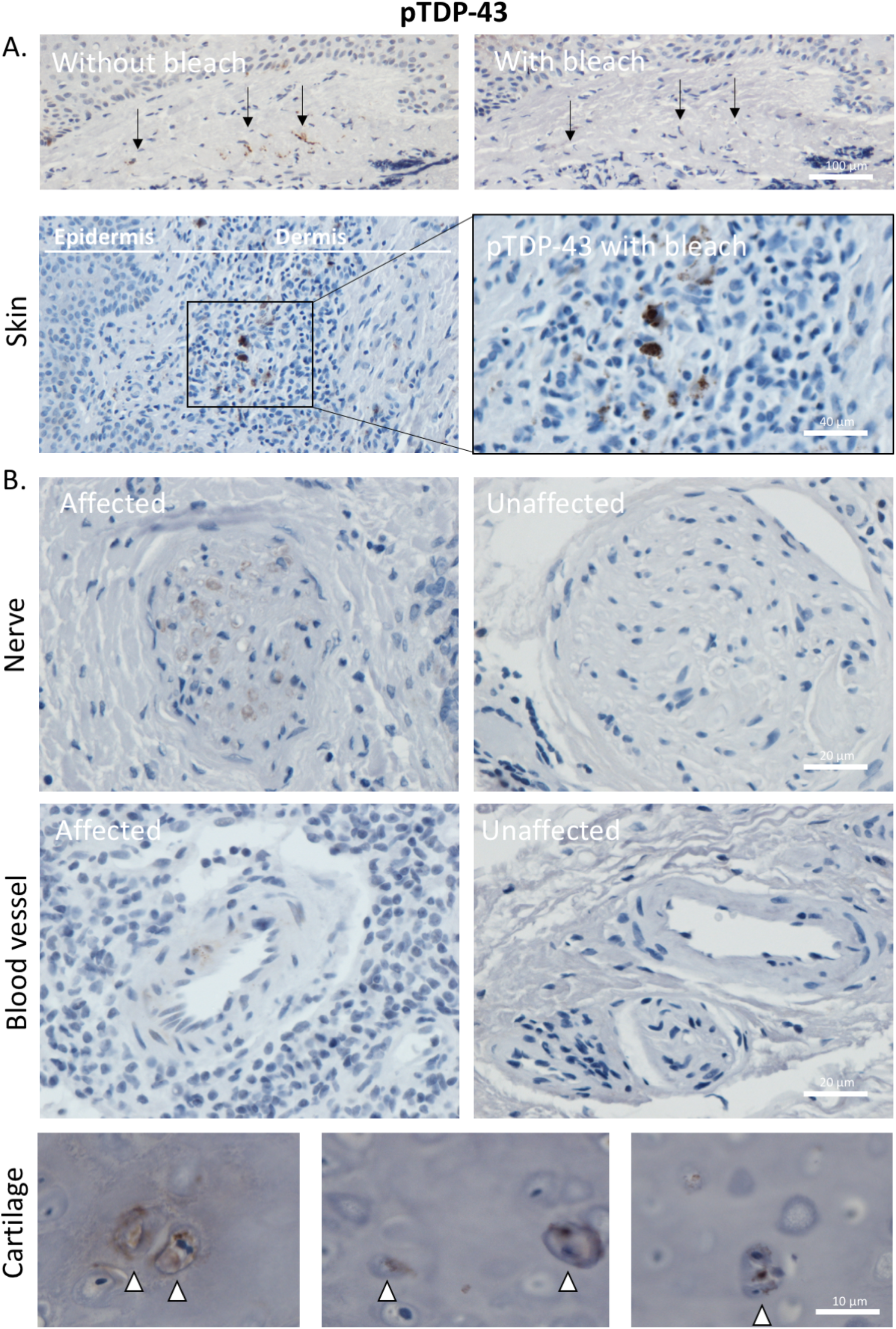
pTDP-43 aggregates identified in nerve bundles, endothelial cells and superficial dermis of skin. **A**. Images demonstrating pTDP-43 staining of skin following removal of brown pigmentation (from melanocytes and/or pigment drop out as a reactive feature) from the dermis and epidermis (above) and pTDP-43 positive aggregates within the superficial dermis (below). All taken from patient 26. **B**. Example of pTDP-43 present within peripheral nerves (top left; patient 14) and an example of an unaffected nerve bundle (top right; patient 29) and within endothelial cells of the dermal blood vessels (bottom left; patient 14) and an unaffected blood vessel for comparison (bottom right; patient 29). **C**. Three example images taken from patient 26 demonstrating pTDP-43 aggregates within the chondrocytes of the ear cartilage sampled as part of an excision specimen for a basal cell carcinoma.

## Discussion

Here we show, for the first time, evidence of pTDP-43 aggregation in human GI tissue taken as part of routine clinical practice during life from ALS patients, prior to diagnosis of their motor symptoms. This finding is in line with gut biopsies taken from patients with Parkinson’s disease, where alpha-synuclein pathology can be seen many years prior to diagnosis [5, 16]. Patients with neurodegenerative diseases frequently report GI symptoms prior to their neurological diagnosis [1]. Our data, as well as that published for alpha-synuclein in PD patients, would support a hypothesis that GI pathology is present before CNS symptoms occur. Interestingly, recently published mouse models of TDP-43 proteinopathy have also shown TDP-43 pathology in the gut. A recent, extensive, phenotypic evaluation of theTDP-43^Q331K^ mouse model stated the mice suffer from gut immotility, due to increased human TDP-43 expression in the myenteric plexus and degeneration of neurons therein, complicating the neurological (CNS) phenotype [10]. This gut phenotype and associated myenteric pathology is also the case for the TDP-43^A315T^ mouse model and the C57BL/6J congenic Prp-TDP43^A315T^ mouse model [11,12]. In all instances, the gut pathology precedes the CNS phenotype and is, in the case of the A315T mouse, too severe causing death of the animal before any indication of CNS disease.

This finding of gut pathology preceding motor symptoms is also seen in Tau, Aβ and alpha-synuclein animal models [17,18]. In these models, it has been hypothesised that the pathology originates in the gut and progresses to involve the CNS when there was (i) a physical connection (intact vagus nerve) and (ii) systemic inflammation [17,18]. When these two conditions were not met, central neuronal pathology did not develop. This raises the question of how pathology arising in the gut might reach the CNS. In our data, and in that published for alpha-synuclein, aggregates are seen in both the lamina propria and the myenteric plexus. Antigens, including soluble molecules of comparable size to TDP-43 and alpha-synuclein, have been shown by two-photon imaging to enter the lumen-facing surface of intestinal goblet cells and be transported to be presented to dendritic cells within the lamina propria [19, 20]. This is referred to as Goblet-cell-associated antigen passages (GAPs) and is restricted to small molecules up to 70 kDa in size. M cells and macrophages are hypothesised to deliver these antigens in this way and may result in a lymphoid reaction, as we see in the gallbladder tissue in our cohort [19, 20]. Alternatively, a physical connection to the CNS has been posited in the animal model literature, the fact that we also detect pTDP-43 aggregates within endothelial cells and in lymph nodes may also raise the possibility of dissemination through a non-physical connection. Furthermore, the fact that we also see pTDP-43 aggregates in chondrocytes of the ear cartilage may also raise the further possibility of meta-synchronous (i.e. occurring separately in a non-linked fashion due to similar environmental stimuli) aggregation occurring in a susceptible individual in certain cell types. Of note, chondrocytes are not unlike motor neurons, as they are long-lived post-mitotic cells, both derived from the neural crest. Therefore, a shared susceptibility to meta-synchronous protein aggregation may be possible.

Whilst the presence of pTDP-43 aggregates within gut tissues in our cohort raises the possibility of a similar mechanism of pathological spread to that seen in other neurodegenerative diseases (as detailed in the early preclinical studies outlined above), we also see pTDP-43 in other tissues including skin, peripheral nerves and blood vessels as well as lymph nodes that would not typically drain from the GI tract. Other groups have identified pTDP-43 aggregates within peripheral nerves, with an identical staining pattern to that seen in our cohort [7]. Several studies have also evaluated TDP-43 pathology in muscle biopsies [8, 9]. One study showed TDP-43 pathology in 19 out of the 57 ALS cases evaluated [9], but a further study showed no evidence of TDP-43 pathology in the three cases that they evaluated [8]. Our cohort only included one muscle specimen; however, we detected no evidence of pTDP-43 pathology in this case. Indeed, these data from other cohorts perhaps indicate that more than just the gut-brain-axis is involved by pTDP-43 aggregation, supporting the hypothesis of meta-synchronous aggregation, rather than physical spreading of pathology. Although the mechanistic consequences of the involvement of other tissues, and their contribution to ALS pathogenesis clearly requires further investigation.

The presence of pTDP-43 aggregates within the blood vessels of 6 out of the 13 cases evaluated in our cohort raises the possibility that TDP-43 pathology could be identified prior to neurological dysfunction in ALS, holding promise as an early detector of pathology. Indeed, endothelial cell involvement by pTDP-43 aggregation could directly result in misfolded forms of TDP-43 in circulating blood, which could be detected in blood samples taken from these individuals prior to symptom onset. Indeed, all cases with endothelial involvement in our cohort had pathology detectable in tissues prior to ALS diagnosis. Furthermore, we have shown previously, by systematic review and meta-analysis, that serum-derived TDP-43 may show promise as a peripheral biomarker in ALS [21].

Our data imply that peripheral non-CNS tissues could hold promise as early indicators of ALS, prior to neurological involvement and that tissue biopsies could also help us to understand how fluid biomarkers, such as blood and stool samples could be utilized through their association with the tissues that they derive from. Early diagnosis, in this way, would not just improve the chances of successful therapeutic intervention, it would also inevitably lead to a rethinking of neurodegenerative conditions from being end-stage neurological disorders to systemic disorders with potential for population screening and early intervention long before neurological manifestations occur.

### Study limitations

Our study was based on a cohort of 48 patients, only 13 of which had surgical specimens that were sufficient for histological assessment, 12 of which were sALS cases. Therefore, the conclusions of this study, whilst informative, are subject to the relevant caveats of a small sample size and are biased towards a predominantly sALS population. For example, with tissues that we have seen no evidence of aggregation in our cohort, we cannot rule out that they are not involved by pTDP-43 aggregation in a larger cohort, we can only state that they are not involved in our cohort. It is therefore important to include all relevant information and detailed methodological reporting to facilitate future meta-analysis where more cases and cohorts can be evaluated in pooled datasets, as has been done for alpha-synuclein [22]. For this reason, we have included detailed clinical information and histological images for future analysis in this way. Another relevant caveat of this work is that we selected only ALS patients who were known to have CNS pTDP-43 pathology. This is therefore a selected cohort and does not allow us to understand population distribution of these pathologies in unaffected individuals. A future study evaluating larger numbers of unselected individuals is now warranted to understand the population dynamics of this pathology and provide more information regarding the feasibility of population screening and the establishment of larger cohorts for molecular studies. However, a great strength of the approach we have taken here, is that surgical biopsies banked in local tissue banks allow us to gain a better understanding of the temporal nature of the development of ALS and the involvement of non-CNS tissues in this process.

## Acknowledgements

The authors would like to thank the staff at the NHS Lothian BioResource (Vishad Patel and Craig Marshall). This study was funded by the Pathological Society (JSPS CLSG 202002 to JMG and JO’S) and the Wellcome trust (108890/Z/15/Z to OR).

